# The Hsp60 C-terminus Senses Substrate and Triggers Allosteric ATP Hydrolysis

**DOI:** 10.1101/2023.09.15.558033

**Authors:** Daniel Von Salzen, Alejandro Rodriguez, Anwar Ullah, Ricardo Andres Bernal

## Abstract

The human mitochondrial chaperonin Hsp60/Hsp10 plays an essential role in maintaining protein homeostasis through an ATP dependent protein refolding mechanism. In the absence of this chaperonin, all cells perish due to an accumulation of misfolded aggregates. Despite its importance, the detailed mechanism of ATP hydrolysis and the role of the C-terminal tail remains unresolved. Here we show that the C-terminal tail acts as a sensor for the arrival of substrate into the chaperonin internal chamber that directly leads to an allosteric trigger of ATP hydrolysis in a neighboring subunit. Our results reveal that removal of the C-terminal tail leads to normal binding of both ATP and a misfolded substrate, but the chaperonin stalls and is unable to progress along the protein folding reaction cycle because it has lost the ability to sense the presence of substrate. High resolution reconstructions reveal the detailed mechanism of ATP hydrolysis where access to the γ-phosphate is blocked by the carboxylate oxygen of an aspartate residue. Binding of the C-terminal tail to substrate displaces this oxygen and allows ATP hydrolysis. These results bring a better understanding of how Hsp60 functions at an atomic level where substrate arrival activates ATP hydrolysis through an allosteric trigger.

## Introduction

Chaperonins play a critical role in the survival of cells by preserving protein homeostasis through the refolding of misfolded proteins (Fan et al., 2020; Fayet, Ziegelhoffer, & Georgopoulos, 1989; Hartman, Hoogenraad, Condron, & Høj, 1992; Hertveldt et al., 2005; Rodriguez, Von Salzen, Holguin, & Bernal, 2020). Eukaryotic cellular survival is dependent on the TRiC/CCT chaperonin for cytosolic protein homeostasis and the mitochondrial heat shock protein 60 (Hsp60) for mitochondrial protein homeostasis (Gestaut et al., 2023; Hildenbrand & Bernal, 2012). Human Hsp60, together with the co-chaperonin Hsp10, carries out this role in the mitochondrial matrix by the folding of nascent and misfolded proteins including those involved in metabolism and electron transport (Cheng et al., 1989; Fan et al., 2020; A. Horwich, 1990; Lubben, Gatenby, Donaldson, Lorimer, & Viitanen, 1990). Monomers of Hsp60 normally assemble into a homo-tetradecameric complex that leads to the formation of an internal protein folding chamber in each of two heptameric rings that are arranged back-to-back(Hildenbrand & Bernal, 2012). These internal chambers sequester misfolded proteins from components of the matrix milieu after binding of the co-chaperonin Hsp10 whose function is to cap the entrance to the internal chambers (Parnas et al., 2012; Vilasi et al., 2014). The resulting conformation of the Hsp60/10 complex is a prolate spheroid that resembles an “American football” where the co-chaperonin forms each pointed end of the complex(Yacob Gomez-Llorente et al., 2020; Nisemblat, Yaniv, Parnas, Frolow, & Azem, 2015). Hsp60/10 substrate refolding proceeds along an ATP-dependent cycle that includes a dramatic ring separation where the back-to-back rings separate during the protein folding stage (Bhatt et al., 2018; Ishida et al., 2018; Levy-Rimler et al., 2001; Molugu et al., 2016; Motojima; Nielsen & Cowan, 1998). Previous studies show that ring separation may allow for the individual internal chambers to increase in volume, providing a more favorable environment to refold particularly large, encapsulated substrates(Bhatt et al., 2018; Brocchieri & Karlin, 2000; Molugu et al., 2016). The single rings come back together after releasing the spent nucleotide and newly refolded substrate, to regenerate the double ring conformation and initiate another protein folding reaction cycle(Jinliang Wang et al., 2019).

An unresolved aspect of the chaperonin reaction cycle is how ATP hydrolysis is triggered to initiate the conformational changes that are required for protein folding(Yacob Gomez-Llorente et al., 2020; Okamoto et al., 2015). While Hsp60 and Hsp10 readily form the prolate spheroid structure in the presence of ATP, the chaperonin will not hydrolyze ATP unless misfolded substrate has been encapsulated(Enriquez et al., 2017). Therefore, there must be a mechanism by which Hsp60 recognizes the presence of substrate in the inner chambers before initiating ATP hydrolysis and subsequent conformational changes.

Within the equatorial domain of Hsp60 exists a small inherently disordered C-terminal tail that is highly conserved among the group I chaperonins(Karlin & Brocchieri, 2000). Only a small portion of the C-terminal tail is ordered and is seen radiating as weak electron density into the internal chamber(Yacob Gomez-Llorente et al., 2020). Its importance in the function of Hsp60 can be seen in yeast where removal of the tail causes complete loss of chaperonin function and eventual cell death(Fang & Cheng, 2002). Some research on the Hsp60 C-terminal tail exists, but most of what is currently known has been inferred from previous studies on the bacterial homolog, GroEL(Brocchieri & Karlin, 2000). The role of the C-terminal tail in GroEL is still highly debated with studies suggesting the tail acts as a substrate shuttle, substrate stabilizer, or even as an active tool in the refolding of misfolded substrate(Chen et al., 2013; Naqvi et al., 2022; Jeremy Weaver et al., 2017; J. Weaver & Rye, 2014). However, other studies have also shown that the C-terminal tail may be coupled to allosteric transitions of bacterial GroEL(McLennan, Girshovich, Lissin, Charters, & Masters, 1993; J. Weaver & Rye, 2014). Here, we have created a series of C-terminal deletion mutants that demonstrate that the Hsp60 C-terminal tail is unnecessary in shuttling misfolded substrate into the chaperonin inner chamber, and instead functions as a sensor that detects the arrival of misfolded substrate in the internal chamber to initiate allosteric ATP hydrolysis and protein folding.

## Results

The aim in this investigation was to determine the role of the Hsp60 C -terminal tail in detecting the arrival of substrate into the internal chamber. Cryo-Electron Microscopy (cryo-EM) is used as a tool to probe the details of not only how the C -terminal tail is able to detect the substrate but how it can then communicate this information to the nucleotide binding pocket for ATP hydrolysis to commence. To explore these questions, we first cloned the Hsp60/10 wild-type chaperonin and a series of deletion mutants to determine the minimal length of C-terminal tail that is required for elimination of ATP hydrolysis while maintaining the ability to assemble the chaperonin.

### C-terminal tail deletion affects Hsp60 assembly and function

A series of C-terminal deletion mutants were generated to determine how much of the tail could be deleted to abolish ATP hydrolysis but retain complex integrity. For simplicity, moving forward we will use the DNA deletion construct nomenclature (i.e., Δ532) to describe the recombinant mutant proteins as well. The first deletion mutant involved removal of the terminal 16 residues (Δ532) to eliminate the M/G rich hydrophobic region (MGAMGGMGGGMGGGMF) (Supplemental Figure S1A). The next mutant involved the removal of 25 residues (Δ523) from the carboxyl-terminus including not only the hydrophobic region but also the proline flanked highly charged (IPKEEKDPG) region. The next two mutants removed 26 and 27 carboxyl-terminal residues (Δ522 and Δ521, respectively) to determine the point at which the chaperonin can still form a tetradecamer but loses its ability to hydrolyze ATP. All the recombinant protein complexes were monitored by DLS and assayed for protein folding activity using denatured MDH as a substrate (Supplemental Figure S1B). DLS measurements indicate that the hydrodynamic diameter of monomeric Hsp60 is about 10nm. When the monomeric Hsp60 is incubated in assembly buffer with ATP, the diameter is increased to ∼17.5 nm confirming the transition from monomer to tetradecameric complex. This hydrodynamic diameter coincides well with previously determined DLS measurements of Hsp60(Vilasi et al., 2014). DLS measurements reveal that most of the purified and assembled recombinant chaperonins have a comparable hydrodynamic diameter to the wild type hsp60 complex of about 17.5 nm (Supplemental Figure S1B). The exception was the Δ521 mutant that resulted in a hydrodynamic diameter of about 10 nm, closely matching the size of the unassembled Hsp60 monomer. The removal of threonine 521 resulted in the instability of β-sheet A composed of strands β1 and β18 of one subunit and β-2 and β-3 of the adjacent symmetry related subunit (full secondary structure description is illustrated in Supplemental Figure S2). Threonine 521 is within β18, and its deletion prevents the formation of the extended β-sheet A between adjacent subunits. Loss of hydrogen bonding provided by T521 destabilizes intersubunit contacts enough to disrupt the entire complex. Since Δ521 does not assemble into a functional chaperonin, it was dropped from further analysis.

The remaining mutant complexes were analyzed for activity to identify the C-terminal residues that might abolish ATP hydrolysis when removed. Each mutant chaperonin was included in a refolding activity assay where denatured human malate dehydrogenase (hMDH), a normal Hsp60 substrate, was used as a substrate in the assay(Dubaquié, Looser, Fünfschilling, Jenö, & Rospert, 1998). Wild-type Hsp60 can refold the denatured hMDH to activity levels equivalent to the native hMDH control (Figure 1A). However, all C-terminal deletion mutants appear to have a diminished denatured hMDH refolding activity. Δ532 was the only deletion mutant able to assist in the refolding of denatured hMDH. Deletion of the hydrophobic poly methionine-glycine tail in Δ532 reduced the activity by about 27% compared to wild-type Hsp60. The fact that about three quarters of the activity remains after removal of the hydrophobic tail indicates it is dispensable to some extent and is not likely involved in substrate capture or active refolding of the substrate itself since a significant amount of activity remains despite its removal. Conversely, with mutant Δ523, deletion of an additional portion of the C-terminal tail that includes the hydrophilic IPKEEKDPG patch abolishes all denatured hMDH refolding activity. Mutant Δ522 has an additional glutamic acid residue deleted and like Δ523 it has also lost all refolding activity. Mutant Δ522 is the last residue that can be deleted before the complex becomes unstable and remains as monomer (Mutant Δ521).

**Figure 1:**
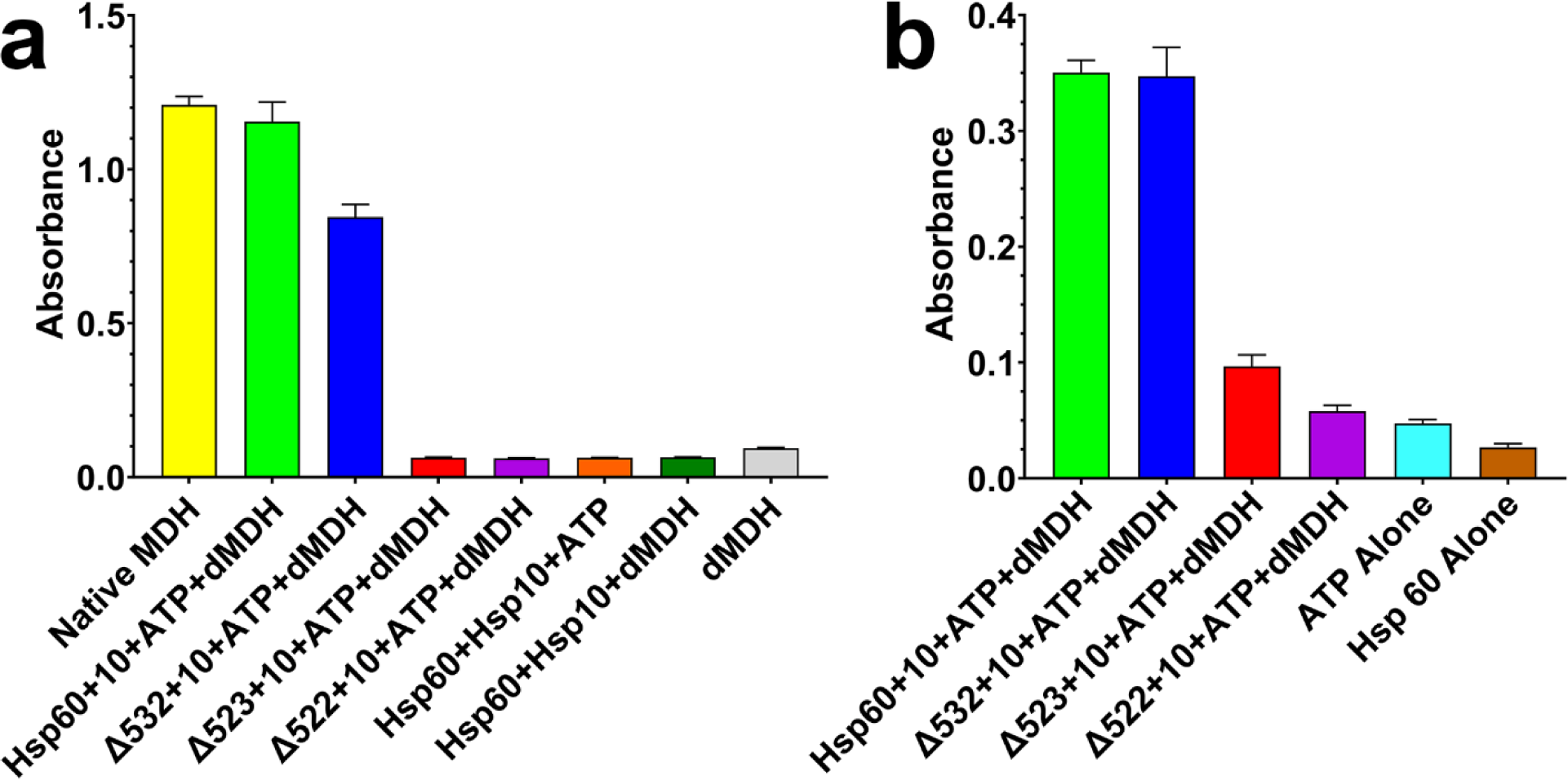
Biochemical characterization of Hsp60/10 recombinant chaperonin and deletion mutants. a) Human malate dehydrogenase (hMDH) refolding assay. The wild-type and C-terminal deletion mutants were assayed for ability to refold denatured hMDH. The assay measures the formation of NADH from NAD^+^ through the conversion of Malate to Oxaloacetate by native hMDH. (b) EnzChek Phosphate Assay to detect ATP hydrolysis. Wild-type and C-terminal deletion mutants were assayed for their ability to refold denatured hMDH. The assay measures the release of inorganic phosphate after ATP hydrolysis. All samples were run in triplicate and error bars represent confidence intervals.

After determining that mutants Δ523 and Δ522 are unable to refold denatured hMDH, we assayed the deletion mutants for their ability to hydrolyze ATP. The idea was to rule out the possibility that the deletion was not affecting other aspects of the protein folding mechanism other than ATP hydrolysis. This assay measures the production of inorganic phosphate, the byproduct of ATP hydrolysis, using the EnzChek phosphate assay kit (Figure 1B). As expected, both Hsp60 and Δ532 showed the ability to hydrolyze ATP, as they both produced high levels of inorganic phosphate. Also of note, Hsp60 in the absence of denatured hMDH does not hydrolyze ATP, consistent with previous studies(Enriquez et al., 2017). However, both Δ523 and Δ522, which were unable to refold denatured hMDH, produced minimal amounts of inorganic phosphate, comparable to normal ATP decomposition levels. This indicated that ATP hydrolysis was severely compromised in Δ523 and Δ522 because of the C-terminal deletions.

### Cryo-EM 2D analysis of wild-type Hsp60 and Δ522

The protein folding assays reveal that Δ522 with 26 residues deleted is a mutant that eliminates protein folding activity and ATP hydrolysis. Furthermore, removal of just one more residue destabilizes the protein such that it is unable to form a tetradecameric complex and remains a monomer. These characteristics made Δ522 the ideal candidate for a high resolution cryo-EM reconstruction because it would reveal how the C-terminal tail is able to affect ATP hydrolysis and protein folding with details at the atomic level. Wild-type Hsp60 and Δ522 were purified and incubated each with the co-chaperonin Hsp10 under the same conditions used for the protein folding activity assays, and in the presence and absence of stoichiometric amounts of denatured hMDH. Several cryo-EM screening micrographs were collected for each of the four samples and 2D class averages were calculated from each (Figure 2A). Wild-type Hsp60 is trapped in the tetradecameric prolate spheroid conformation without substrate (Figure 2A top). When substrate is added, wild-type Hsp60 progresses along the reaction cycle to fold a protein and the 2D class average obtained is the tetradecamer without Hsp10 in a conformation we normally recognize as the APO conformation where the nucleotide binding pocket is empty. In this sample, the wild-type chaperonin was able to detect the presence of substrate, which then induced ATP hydrolysis and normal reaction cycle progression. The progression of the chaperonin from the prolate spheroid particles to the open double ring conformation with the addition of denatured substrate is in agreement with previous EM observations(Ishida et al., 2018). Removal of the C-terminal 26 residues in Δ522 abolished the ability of the chaperonin to detect the presence of substrate and the 2D class averages for both samples with and without substrate resulted in the tetradecameric prolate spheroid conformation (Figure 2A bottom). Even in the presence of substrate, ATP hydrolysis is not triggered. An interesting observation is that the addition of substrate seems to stabilize the prolate spheroid conformation as many more complexes were identified in those micrographs compared to the micrographs of the sample without substrate that had a noisier background with broken particles.

**Figure 2:**
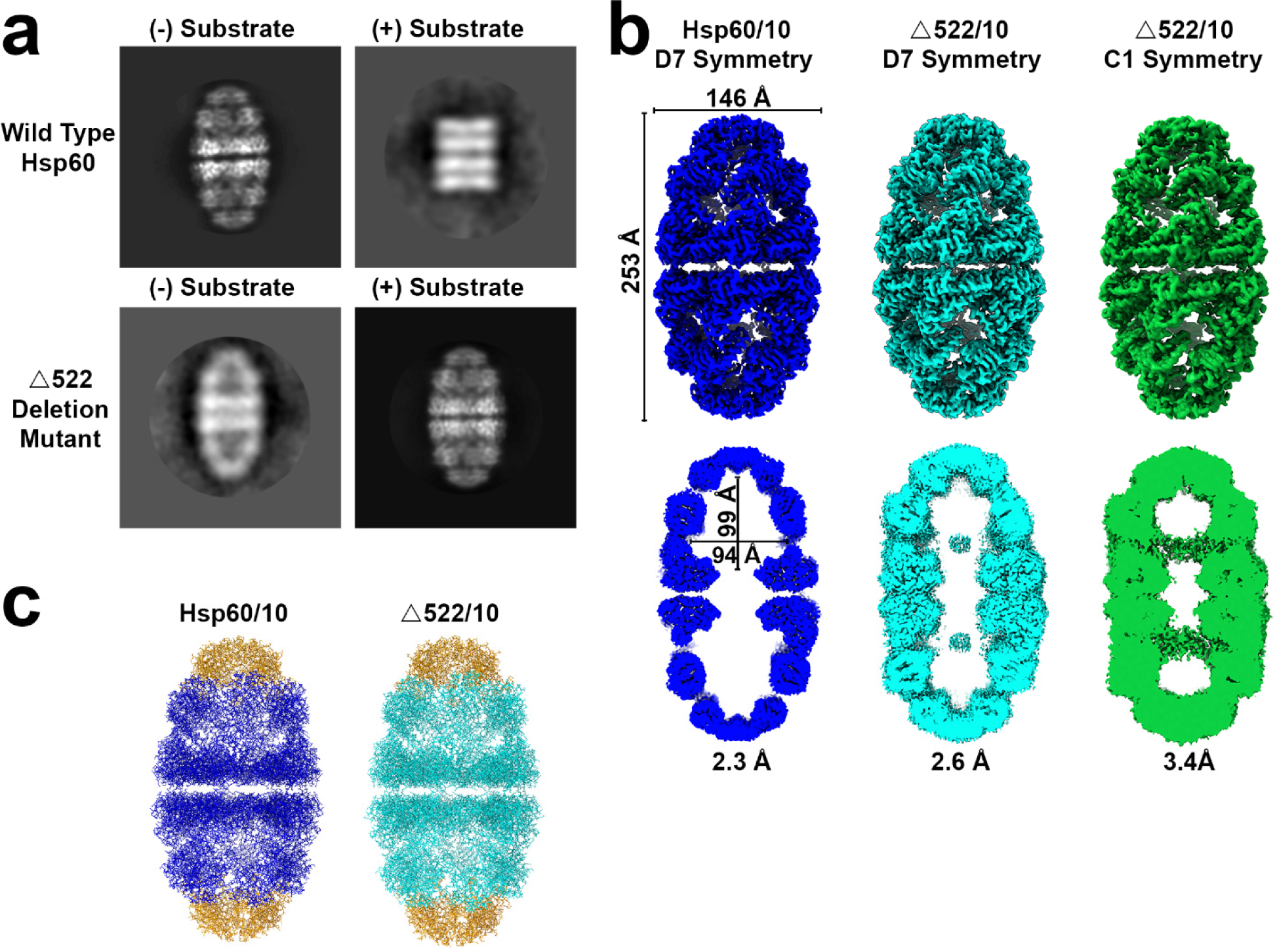
Cryo-EM structural characterization of Hsp60/10 recombinant chaperonin and deletion mutant Δ522. a). 2D Class averages of Hsp60/10 and Δ522 in the presence and absence of denatured substrate to determine if ATP is being hydrolyzed. b). Cryo-EM reconstructions of Hsp60/10 wild-type (**Blue**), Δ522/10 (**Cyan**), and Δ522/10 (**Green**) without symmetry imposed. Below each reconstruction is a slab view to illustrate the of the internal chamber. The Δ522 reconstruction in cyan and green were processed with D7 or C1 symmetry, respectively to see if more of the substrate became ordered. Substrate is visible in both rings implying that Hsp60 folds a protein in each of the two protein folding chambers simultaneously. c). Atomic coordinates were built into corresponding reconstructions in panel B.

### Cryo-EM Reconstructions

Cryo-EM data for the human Hsp60/Hsp10 wild-type chaperonin without substrate was collected at the National Cryo-Electron Microscopy Facility (NCEF) in Frederick, Maryland. We obtained a reconstruction of the tetradecameric prolate spheroid (so-called “football” conformation) to 2.3 Å resolution in the presence of an equimolar amount of Hsp10 and 2mM ATP (Figure 2B, Blue). The raw micrographs, 2D class averages, and the final reconstruction all reveal a single conformation with ATP still bound to the nucleotide binding pocket. This wild-type chaperonin has external dimensions of 253 Å nm from the tip of the Hsp10 co-chaperonin to the other and a width of 146 Å at the widest part of the equatorial region. This is almost 10 Å longer than the published 3.15 Å X-ray crystal structure (PDB 4PJ1). The inner chamber has dimensions of 99 x 94 Å (Figure 2B, blue).

Cryo-EM data for the Δ522 deletion mutant was also collected at the National Cryo-Electron Microscopy Facility (NCEF) in Frederick, Maryland. This data yielded a reconstruction to 2.6Å resolution. This sample was prepared with Hsp60, Hsp10, ATP/Mg^+2^, and included denatured hMDH as substrate (Figure 2B, Cyan). The substrate is visible in the center of the internal chamber but is completely disordered indicating that the refolding process has not commenced. The reconstruction was repeated with the symmetry relaxed to see if more of the substrate was ordered, but this reconstruction too confirmed that the refolding process has not started, and the substrate is completely disordered with only random density appearing (Figure 2B, green). Δ522 chaperonin has the same dimensions as the wild-type structure. The built atomic coordinates have an r.m.s.d. of about 0.35 Å between wild-type and Δ522, indicating that the structures are the same except for the deleted 26 C-terminal residues in the Δ522 mutant.

Atomic coordinates for both reconstructions were modeled into the cryo-EM reconstructions using the program COOT and refined using the program Phenix (Supplemental Table S1). Refinement resulted in excellent geometries with no residues in the Ramachandran disallowed region. Cryo-EM reconstruction maps and model coordinates were deposited into the wwPDB OneDep System with the following accession codes; Wild type PDB 8U4M and EMD-41887, as well as mutant Δ522 PDB 8U4G and EMD-41881.

### Equatorial Contacts in the Double Ring Complex

A distinguishing feature of the Hsp60/Hsp10 chaperonin is that it forms single rings as part of the normal protein folding reaction cycle(Bhatt et al., 2018; Y. Gomez-Llorente et al., 2020; Molugu et al., 2016; J. Wang et al., 2019). The first crystal structure of Hsp60 to 3.2 Å resolution (PDB 4PJ1) was reported to have equatorial asymmetries that protruded away from the chaperonin(Nisemblat et al., 2015). The reconstruction presented here does not have any of the equatorial asymmetries seen in that crystal structure and instead has perfect D7 symmetry in agreement with the previously published cryo-EM structure(Y. Gomez-Llorente et al., 2020). Even after the Δ522 reconstruction was processed without symmetry (C1), the resulting reconstruction to 3.4 Å clearly still had D7 symmetry and the coordinates from the D7 reconstruction fit perfectly without the need for adjustment. Our wild-type and Δ522 reconstructions reveal an equatorial gap between the rings where most residues are no closer than 6-11 Å between rings (Figure 3). Each subunit has a modest contact region between the two rings that is composed of hydrophobic symmetry related residues L463 and E466 as well as a hydrogen bond between dihedral symmetry related serine 462 residues (Figure 3).

**Figure 3.**
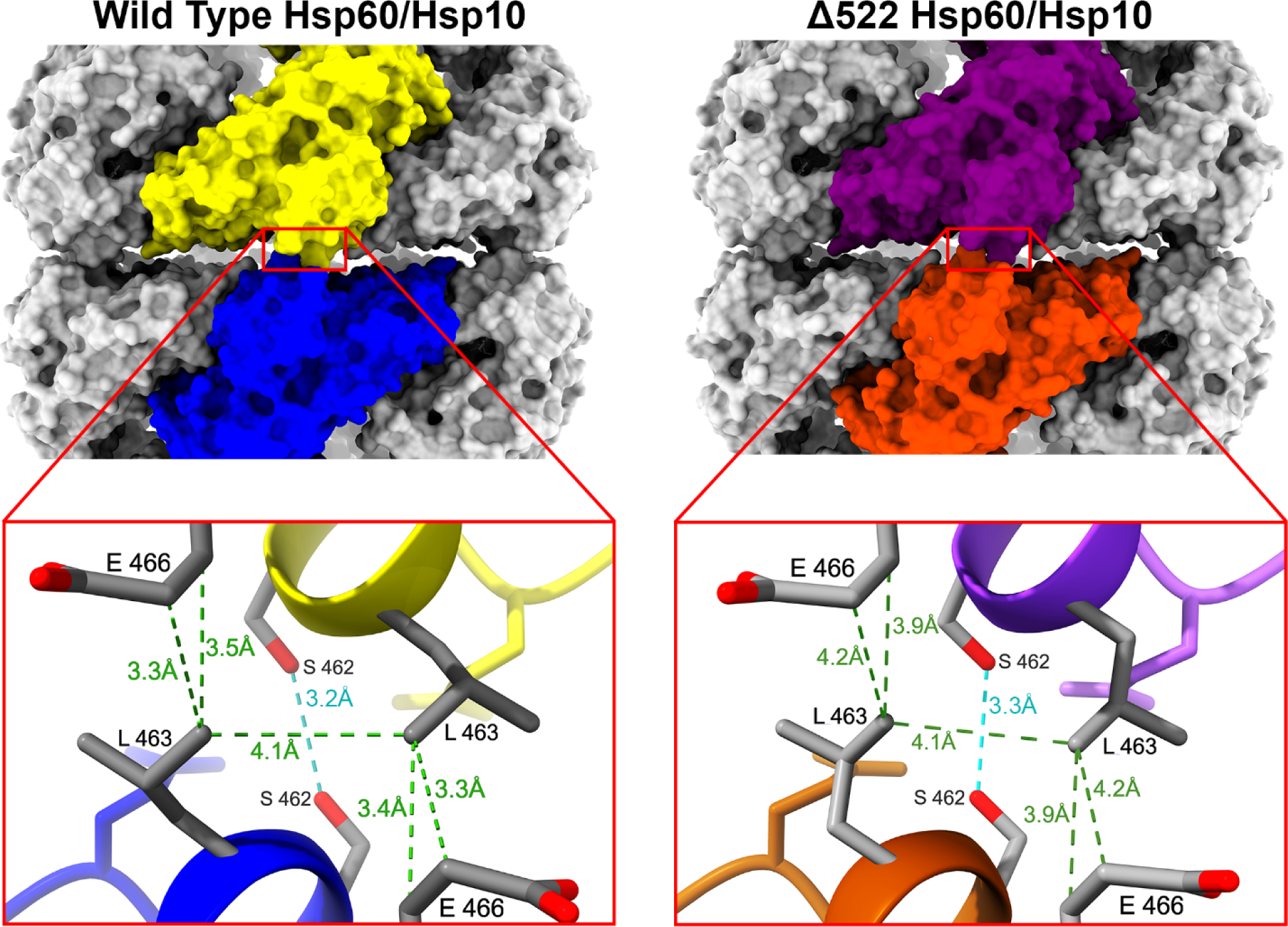
Inter-ring contacts in the equatorial domain for both wild-type Hsp60 and Δ522 prolate spheroid chaperonins. Inset regions boxed in red reveal a closer look at one of the seven contact regions that hold the two rings together. The **green** pseudo-bonds represent Van der Waals contacts while a hydrogen bond is designated **cyan**.

### Nucleotide Binding Pocket and Ligands

The nucleotide binding pocket in both reconstructions is well ordered with the Δ522 reconstruction revealing remarkable details in what appears to be a nucleotide binding site poised for ATP hydrolysis. The triphosphate end of the ATP molecule is coordinated to two cations that we believe are magnesium ions responsible for lining up the ATP for gamma phosphate hydrolysis (Figure 4). The first Mg^2+^ is coordinated in a tridentate configuration to oxygens from each of the three ATP phosphate groups while a second Mg^2+^ is found on the opposite side of ATP and is coordinated in a bidentate configuration to oxygens on the alpha and gamma phosphates of ATP(Boisvert, Wang, Otwinowski, Norwich, & Sigler, 1996; Buelens, Leonov, de Groot, & Grubmüller, 2021). We identify both cations as Mg^2+^ because we stripped the nucleotide binding pocket of both nucleotide and cations before running our sample on size exclusion chromatography to produce an APO chaperonin. It is only after producing the APO conformation that we added ATP and Mg^2+^. The bidentate cation is identified in the literature as a K^+^ ion, but we stress that the identity of this second cation is only assumed to be a potassium(Chaudhry et al., 2003). Positive identification of the bidentate cation would require synchrotron X-ray data collection utilizing the cation absorption edges or neutron diffraction, but this is outside the scope of this study. For our purposes, all that is important is that there are two cations coordinating with the ATP to orient the nucleotide for subsequent hydrolysis of the γ-phosphate. The Mg^2+^ cations are both coordinated with an octahedral geometry as reported previously in the literature (Figure 4) (Pontikis, Borden, Martínek, & Florián, 2009). The tridentate Mg^2+^ is coordinated to an oxygen on each of the three phosphates of ATP, a carboxylate oxygen on residue D85, and two water molecules. The bidentate Mg^2+^ is coordinated to the alpha and gamma phosphate oxygens on ATP, one water molecule, a main chain carbonyl oxygen and side chain gamma oxygen of T28 as well as the main chain carbonyl oxygen of K49. It has been suggested that perhaps substrate binds to the Hsp60 chaperonin before ATP but our 2.3 Å reconstruction of Hsp60 in the presence of ATP and no substrate, clearly demonstrates that ATP has bound in the absence of substrate (Figure 4)(Mas et al., 2018). Of note on this 2.3 Å reconstruction is the weaker electron density for the bidentate Mg^2+^ cation indicating a possible lower occupancy.

**Figure 4.**
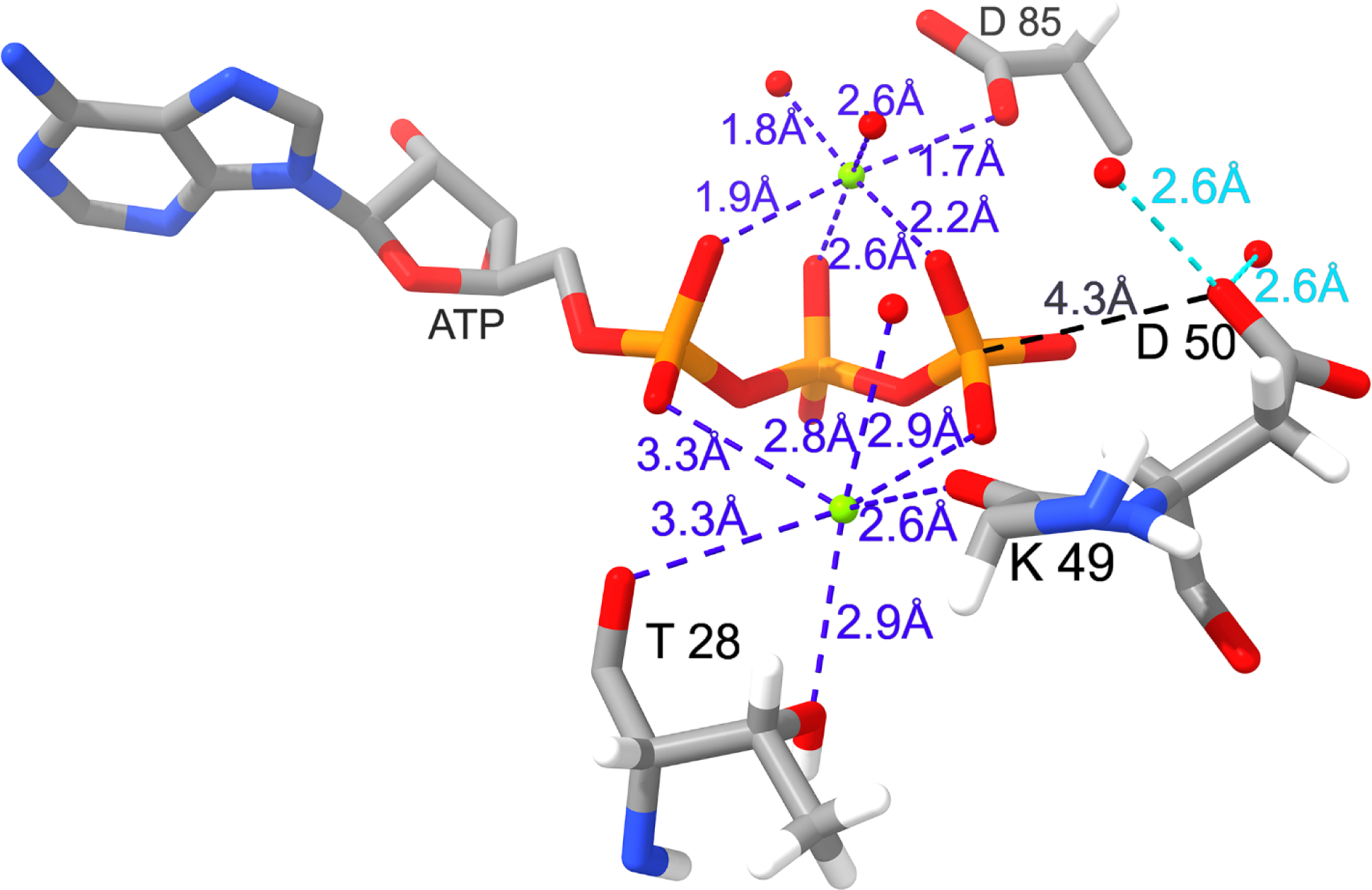
ATP interactions with residues or ligands in the ATP binding pocket of Δ522. Two **Mg^2+^** cations were built into the structure at the locations designated by the green sphere. Coordination atoms were then determined based on distance as indicated in the figure. Each cation has octahedral coordination and includes ordered water molecules (designated as the red spheres).

### C-terminal Truncation

The C-terminal deletion mutation involved removing residues that are normally disordered and so we did not anticipate structural changes to the chaperonin. The ATP binding site of the Δ522 compared to the WT Hsp60 reveals that the two chaperonins remained relatively unchanged even with the deletion of the C-terminal tail in Δ522, indicating that Δ522 should still be capable of ATP hydrolysis(Balasubramanian & Stitt, 2010; Thomsen & Berger, 2008). The activity assays in Figure 1, however, clearly indicate that the loss of the C-terminal tail affects ATP hydrolysis and so we next looked for a structural explanation for these observations.

The part of the C-terminal tail that is still ordered is seen coming away from the wall of the interior cavity towards the direction of where the substrate is expected to be located. The 26 truncated residues would therefore continue to extend out into the interior cavity as indicated in Figure 5. Hsp60 subunits located next to each other in each of the rings interact to form β sheet A with two strands from each adjacent subunit. The amino terminus forms strand β-1 which interacts with the carboxy terminal strand β-18 from the same subunit (Figure 5A). Each Hsp60 subunit contributes the β-2 and β-3 strands (β-2/β-3 loop) that then hydrogen bond to the terminal strands β1 and β18 of the neighboring subunit to form the four stranded β sheet A (Figure 5A, green/blue β sheet). The process is repeated for a total of 7 β sheets in each ring. Interestingly, β-3 ends near residue D50 where the D50-Oδ1 carboxylate oxygen is remarkably close to the γ-phosphate of the ATP, providing an indication of how premature ATP hydrolysis is prevented.

**Figure 5:**
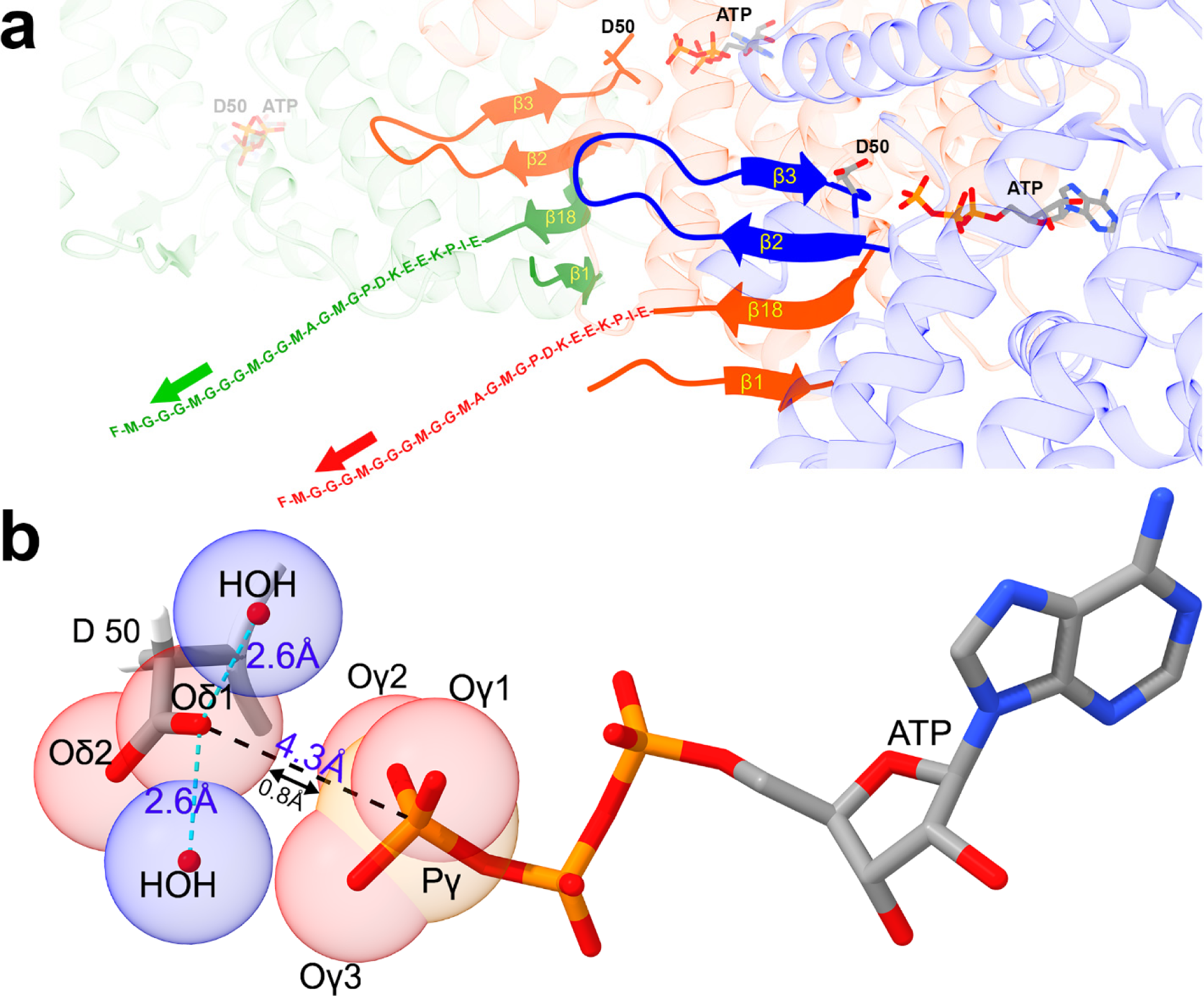
Relationship between the Hsp60 C-terminal tail to the ATP binding site. a). Three C7 symmetry related subunits (green, red, and blue) oriented to view them from the inner chamber reveals the composition of β sheet A. This β sheet is composed of strands **β1**/**β18** from one subunit and stem loop strands **β2/β3** of the neighboring subunit (blue). b). An enlarged view of residue D50 and its proximity to the γ-phosphate of ATP. Relevant atoms have been converted to space filling atoms to better appreciate the limited space in the area. There are two water molecules hydrogen bonded to the D50-Oδ1 and are represented by a blue sphere.

When we look closer at the conserved aspartate D50-Oδ1 in the reconstruction, there is slightly weaker density located at either side of this carboxylate oxygen, which we attributed to ordered water molecules (Figure 5B, blue spheres). When waters molecules were placed there, the distance from the carboxylate oxygen was 2.6 Å for each water, indicating that they were both hydrogen-bonded to the carboxylate oxygen. Furthermore, when the D50 carboxylate oxygens, the water molecules, and the γ-phosphate atoms were depicted as space-filling spheres, it became evident that the D50-Oδ1 was preventing the water molecules from approaching the γ-phosphate thereby preventing premature γ-phosphate hydrolysis (Figure 5B). The distance between the D50-Oδ1 and the γ-phosphate is 4.3 Å from center to center. If the radius of an oxygen is 1.52 Å and the radius of a phosphorus is 1.95 Å then the gap remaining between the two atoms is only about 0.8 Å.

## Discussion

In the work presented here, we used single particle cryo-Electron Microscopy and biochemical assays to characterize the function of the Hsp60 C-terminal tail. What started as a simple question has revealed that the detection of substrate entering the chaperonin triggers allosteric ATP hydrolysis in a neighboring chaperonin subunit. The two reconstructions contained herein reveal an identical structure with strong density for bound ATP. The only difference was the absence of substrate and weaker density for the bidentate Mg^2+^ in the 2.3 Å resolution reconstruction. Our working hypothesis is that the substrate is recruited through hydrophobic interactions between the apical domain and misfolded substrate that leads to internalization into the cavity by additional interactions with the apical domains and interior chamber residues. As the denatured substrate enters the internal chamber, it interacts with the C-terminal tail of the Hsp60 subunits. This interaction tugs at the C-terminal tail towards the inner chamber. The C-terminal tail near the wall of the internal cavity forms a beta strand (β18) that is hydrogen bonded to β1 of the amino terminus, and to β2 and β3 of a neighboring subunit to form the four strand β-sheet A (Figure 5A). At the end of the β3 strand, is aspartate 50 which lies up against the γ-phosphate of ATP. Carboxylate oxygen (δ1) of D50 is located adjacent to the γ-phosphate leaving only a 0.8 Å gap between the two atoms (Figure 5B). This arrangement effectively blocks access to the γ-phosphate and prevents premature γ-phosphate hydrolysis before substrate enters the internal protein folding chamber and blocks uncontrolled hydrolysis of ATP in non-productive chaperonin reaction cycles. Next, hydrolysis of ATP requires the retraction of this D50-Oδ1 away from the γ-phosphate of ATP to allow a water molecule access to the γ-phosphate for a nucleophilic attack to take place. The D50-Oδ1 retraction occurs when the misfolded substrate interacts with the C-terminal tail (Figure 6). Additionally, surrounding the D50-Oδ1, are two ordered water molecules that play a critical role after retraction of D50-Oδ1 (Figure 5B). These two water molecules are hydrogen bonded to the D50-Oδ1 and as the retraction of D50 occurs, one of the water molecules (each would have an equal chance) slides into the space between the D50-Oδ1 and the γ-phosphate (Figure 6). At this point, the conserved aspartate D50-Oδ1 transiently accepts a proton from a hydrogen bonded water and thus acts as catalytic base. The oxygen from the water then executes a nucleophilic attack on the γ-phosphate of the ATP molecule. In a second step, the abstracted proton is transferred to the second water that is hydrogen bonded to the same D50-Oδ1. This water then leaves as a hydronium ion (Figure 6). A study on ATP hydrolysis in the ABC transporter suggests that the abstracted proton is transferred to the inorganic phosphate to form H_2_PO ^-^ instead of HPO ^2-^ after the nucleophilic attack has occurred(Priess, Goddeke, Groenhof, & Schafer, 2018). However, in the Hsp60 chaperonin, the movement that retracted D50 away from the gamma phosphate means that such a mechanism would necessitate that the proton traverses a span of more than 3 Å to reach the HPO_4_ ^2-^. Donating the proton to a water that is already hydrogen bonded to the D50-Oδ1 makes more mechanistic sense(Afanasyeva et al., 2014; Parke, Wojcik, Kim, & Worthylake, 2010).

**Figure 6:**
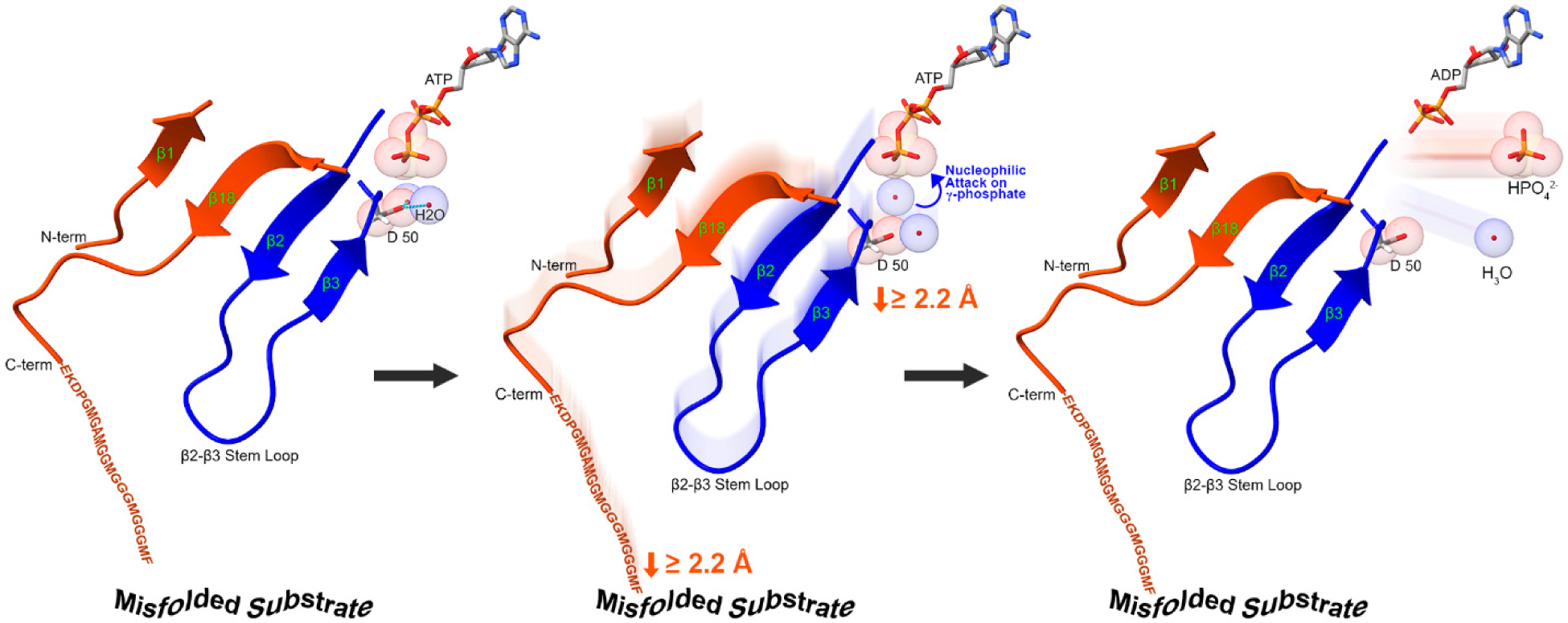
Mechanism of ATP hydrolysis involving the C-terminal tail and the Oδ1 carboxylate oxygen on aspartate 50. β sheet A, relative to the two subunits (orange and blue), has been reoriented compared to Figure 5a to better illustrate the ATP hydrolysis mechanism. Binding of the C-terminal tail to the substrate will tug at β sheet A which will in turn move the D50 residue away from the ATP γ-phosphate. One of the ordered water molecules will slip into the space between D50-Oδ1 and the ATP γ-phosphate. Immediately, the D50-Oδ1 will transiently abstract a proton from this water and the hydroxyl will then execute a nucleophilic attack on the ATP γ-phosphate. The γ-phosphate is converted to HPO ^2-^ and the abstracted proton located on the D50-Oδ1 is donated to the second hydrogen bonded water molecule which is then released as a hydronium ion.

From the onset, our studies sought to replicate native structures and biological ligands to avoid the possibility of cryo-EM reconstruction with artifacts. Therefore, other than deleting the C-terminal tail, our proteins were constructed to mimic native proteins found *in-vivo* without adding mutations that might stabilize the protein or complex for structural studies. We found that the protein complexes were in fact fragile and now that we know that the inter-ring contacts are so minimal, we can understand the reason behind that fragility. In fact, chaperonin samples that are treated harshly, such as with vigorous pipetting, result in a high percentage of single rings. We refer to these single rings as off pathway intermediates because we think they are the result of harsh treatment of the sample and are not a legitimate intermediate during protein folding. We carried the same philosophy with the use of ligands. We used only natural nucleotides instead of non-hydrolyzable analogs to mimic the *in-vivo* complex as closely as possible.

The Hsp60 ATP hydrolysis mechanism presented here is supported by strong biochemical data. Wild-type Hsp60 and Hsp10 in the presence of ATP, stalled in the absence of misfolded protein substrate resulting in our 2.3 Å reconstruction. As soon as substrate was added to this same preparation, a mixture of conformational states was obtained upon ATP hydrolysis. Removal of the C-terminal 26 residues (mutant Δ522) resulted in a chaperonin that not only stalled in the absence of substrate but failed to hydrolyze ATP even in the presence of substrate. This strongly indicates that the C-terminal tail participates in not only sensing the presence of substrate but also in initiating ATP hydrolysis once substrate is detected. Considering the amino acid identity between human Hsp60 and *E. coli* GroEL is 50.9%, we suggest the same allosteric ATP hydrolysis mechanism is utilized in the GroEL/GroES chaperonin.

## Conclusion

These results describe how Hsp60 functions at an atomic level where substrate arrival activates ATP hydrolysis through an allosteric trigger. Our high-resolution reconstructions clearly demonstrate the position of critical water molecules poised to move into a transiently blocked location between the γ-phosphate of ATP and the carboxylate oxygen of the catalytic residue D50. Movement of D50 away from ATP to give the water molecule access to the γ-phosphate is triggered by substrate arrival and binding to the C-terminal tail of a symmetry related and adjacent Hsp60 subunit.

## Methods

### Chaperonin Cloning and Expression

All Hsp60 clones used in this study included removal of the N-terminal mitochondrial targeting sequence. Truncated constructs of the HSPD1 (Hsp60) gene were first generated by removing 16, 25, and 26 residues from the C-Terminal end through PCR and cloned into the pET-11d expression vector between the NcoI and BamHI restriction sites. Transformed BL21 cells were grown to an optical density of 0.6 before inducing with 0.1 mM Isopropyl β-D-1-thiogalactopyranoside (IPTG) for 3 hours at 30°C(Nisemblat, Parnas, Yaniv, Azem, & Frolow, 2014). The cells were centrifuged at 4,000 x g for 30 min at 4°C and were resuspended (1:3 w/v) in a hypotonic buffer that contained 25 mM TRIS base pH 7.5, 50 mM EDTA, 50 mM NaCl and 1 mg/mL of lysozyme. The cells were subjected to 3 rounds of freeze/thaw cycles. DNase I was added at 0.1 mg/mL along with 100 mM MgCl_2_. Hsp60 and each deletion mutant were treated with 40% ammonium sulfate and pelleted by centrifugation at 10,000 x g for 30 minutes. The pellet was resuspended (1:2 w/v) in chromatography buffer containing 25 mM TRIS base pH 7.5 and 10 mM EDTA.

### Purification and Assembly of Hsp60 WT and CTM Mutants

The ammonium sulfate treated Hsp60, and deletion mutant preps were first diluted tenfold with chromatography buffer before being loaded through a Hi Trap Q XL column (GE Healthcare). A linear gradient of sodium chloride from 0 to 1 M was used to purify the chaperonins where each eluted at a conductivity of 5.5 mS/cm. Fractions containing the monomeric Hsp60 or mutant protein were pooled and concentrated through a Vivaspin 20 MWCO 10,000 spin column before treatment with a reassembly buffer containing 25 mM TRIS pH 7.5, 12 mM KCl, 12 mM MgAcetate, and 4 mM ATP at 27°C for one hour to assemble the chaperonin complexes. After assembly, the sample was treated with 100 mM phosphate buffer pH 7.5 and 50 mM EDTA before purification by gel filtration at room temperature on a Superose-6 Increase column (GE Healthcare) equilibrated with buffer containing 25 mM TRIS pH 8, 150 mM NaCl, 12 mM MgCl_2_(A. L. Horwich & Fenton, 2009; Viitanen et al., 1998). Both Hsp60 and deletion mutant complexes were collected and concentrated using a Vivaspin 20 MWCO 10,000 spin column. After concentration, wild-type Hsp60 was flash frozen using liquid nitrogen for storage while all deletion mutant samples were assessed the same day after purification. Protein purification was confirmed by sodium dodecyl sulfate polyacrylamide gel electrophoresis (SDS-PAGE) with a single band at ∼60 kDa.

### Expression and Purification of Hsp10

The HspE1 (Hsp10) gene was cloned and inserted into the pET-22b (+) expression vector at the NdeI and XhoI restriction sites and transformed into *E. Coli* BL21 (DE3) cells. The BL21 cells were grown to an optical density of 0.6 before inducing with 0.1 mM IPTG for 3 hours at 30°C. The cells were centrifuged at 4,000 x g for 30 min at 4°C and were resuspended (1:3 w/v) in a hypotonic buffer that contained 25 mM TRIS base pH 6.5, 50 mM EDTA, and 50 mM NaCl. The resuspended cells underwent the same freeze-thaw treatment as was done with Hsp60. The protein pellet was finally resuspended (1:2 w/v) in chromatography buffer containing 25 mM TRIS base pH 6.5, 50 mM NaCl, and 10 mM EDTA.

The ammonium sulfate treated Hsp10 preps were first diluted tenfold with chromatography buffer before loading into a HiPrep SP XL 16/10 column (GE Healthcare). A sodium chloride linear gradient from 0 to 1 M was used to purify Hsp10 and it eluted at a conductivity of 13.2 mS/cm. Hsp10 fractions were pooled and concentrated using a Vivaspin 20 MWCO 10,000 spin column before purification on a Superose-6 Increase column (GE Healthcare) equilibrated with buffer containing 25 mM TRIS pH 8, 150 mM NaCl, and 12 mM MgCl_2_. Hsp10 fractions were collected and concentrated using a Vivaspin 20 MWCO 10,000 spin column before being flash-frozen with liquid nitrogen for storage.

### Expression and Lysis of Human Malate Dehydrogenase

The human MDH2 (hMDH) gene was inserted into the pET-22b (+) expression vector at the NdeI and XhoI restriction sites and transformed into *E. Coli* BL21 (DE3) cells. The BL21 cells were grown to an optical density of 0.9 before inducing with 2 mM IPTG for 1 hour at 30°C. The cells were centrifuged at 4,000 x g for 30 min at 4°C and were resuspended (1:3 w/v) in a hypotonic buffer that contained 25 mM TRIS base pH 7.5, 50 mM EDTA, and 50 mM NaCl. The resuspended cells underwent the same freeze/thaw treatment as was done with Hsp60. The DNase I treatment and centrifugation, the supernatant was treated with ammonium sulfate precipitate protein between the 40-60% range. The final protein pellet was dissolved (1:2 w/v) in chromatography buffer containing 25 mM TRIS base pH 7.5, 1.5 M ammonium sulfate, and 10 mM EDTA.

The ammonium sulfate treated hMDH preps were loaded onto a 5 mL HiTrap Butyl HP column (GE Healthcare). An inverse linear gradient of ammonium sulfate from 1.5 M to 0 M was used to purify hMDH where it eluted at a conductivity of 135 mS/cm. hMDH fractions were first buffer exchanged into buffer containing 25 mM TRIS pH 8, 150 mM NaCl, and 12 mM MgCl_2_ before being pooled and concentrated using a Vivaspin 20 MWCO 10,000 spin column. The concentrated sample was then purified by gel filtration on a Superose-6 Increase column (GE Healthcare) equilibrated with the same buffer used for the buffer exchange. hMDH fractions were collected and concentrated before being flash-frozen with liquid nitrogen for storage.

### Dynamic Light Scattering

After purification, complex formation of Hsp60 and all deletion mutants were examined by dynamic light scattering using a Malvern Zetasizer Nano ZS to ensure the complexes were formed. Protein samples were diluted to final concentration of 0.5 mg/mL in buffer containing 25 mM TRIS pH 8, 150 mM NaCl, 12 mM MgCl_2_. These samples were read at 37°C and all data was collected in triplicates.

### Activity of C-terminal Mutant Proteins

The chaperonin activity of all deletion mutants was analyzed through the refolding of the substrate hMDH. hMDH was first denatured with 20 mM HCl and 1 mM DTT to a final concentration of 62.5 nM and left to incubate at room temperature for 20 min. After incubation, MDH was neutralized with 50 mM TRIS pH 7.5. An assay mixture was prepared that contained 6.5 μM of the denatured MDH, 91μM of an Hsp60 mutant chaperonin, 91 μM of Hsp10, and 2 mM ATP in the same buffer used for size exclusion chromatography. The protocol outlined in the Malate Dehydrogenase Assay Kit by Sigma was followed to measure the renaturation of denatured hMDH using each chaperonin preparation.

The ATP hydrolysis activity of each chaperonin preparation was evaluated using the EnzChek Phosphate Assay Kit. hMDH was denatured similarly as was done in the activity assay. The denatured hMDH was then incubated at 22°C with 200 μM MESG, 0.2 units Purine Nucleoside Phosphorylase (PNPase), 2 mM ATP and the kit’s reaction buffer for 10 min. After incubation, 91 μM of Hsp60 and 91 μM of Hsp10 were added and a continuous colorimetric analysis was immediately started. The absorbance was measured at 360nm for 30 min at 22°C.

### Electron Microscopy

The ATP conformation of the Hsp60 wild type and the Hsp60 deletion mutant in the presence of denatured substrate were prepared for cryo-electron microscopy in a very similar way. The samples were prepared by mixing Hsp60 with Hsp10 in an equimolar ratio. The Δ522 was also mixed with denatured hMDH substrate in a 7:1 chaperonin to substrate ratio. Both chaperonin samples were placed in buffer containing 25 mM TRIS pH 8, 12 mM MgCl_2_, 4 mM ATP, to a final concentration of 3 mg/mL of protein. Three microliters of sample were applied onto a glow discharged 300-mesh Quantifoil R1.2/1.3 holey carbon grid. The excess liquid was blotted off for 7 seconds from the carbon side of the grid before plunging it into liquid ethane using a manual plunger. The flash frozen samples were imaged on a 300 kV Titan Krios microscope equipped with a Gatan K3 camera at the National Cryo-EM Facility (NCEF) in Frederick Maryland(Baxa et al., 2020). Data collection parameters are summarized in Supplemental Table S1.

### Image Processing and Structure Determination

Both reconstructions were processed using the RELION software package(Scheres, 2012). All movies were motion corrected using the RELION motion correction implementation. CTF estimation was done from RELION using CTFFIND4(Rohou & Grigorieff, 2015). Several hundred particles were manually picked and used to calculate 2D class averages. Class averages with side and top views were used to automatically pick 586,721 wild type particles from 6400 micrographs and 1,145,473 Δ522 mutant particles from 7929 micrographs. The particles were extracted using a box size of 320 x 320 pixels before being subjected to multiple iterations of 2D classification. Bad 2D class averages and the particles within each bad average were removed from the dataset after each iteration until most of the bad particles were deleted. After an initial 3D refinement, a 20 Å lowpass filtered mask was created in addition to running RELION CTF refinement. This resulted in a reconstruction of 2.9 Å for the wild type sample and 3.0 Å for the Δ522 mutant. The resolution increased to 2.3 Å for the wild type sample and 2.6 Å for the Δ522 mutant based on a Fourier shell correlation threshold of 0.143 after postprocessing that included applying a B-factor correction of -124.4 to the wild type reconstruction and -121.8 to the Δ522 mutant. Coordinates were then modeled into the reconstructions using Coot and refined using the program Phenix(Afonine et al., 2018; Emsley, Lohkamp, Scott, & Cowtan, 2010). An additional 3D-refinement cycle was run on the Δ522 mutant with C1 symmetry instead of D7 to generate the no symmetry reconstruction in Figure 2.

## Data Availability

Cryo-EM reconstruction maps and model coordinates were deposited into the wwPDB OneDep System with the following accession codes; Wild type PDB 8U4M and EMD-41887, as well as mutant Δ522 PDB 8U4G and EMD-41881. Raw cryo-EM micrograph datasets are deposited into the EMPIAR database.

## Author Contributions

D.V.S. performed most experiments, A.R. helped with some of the experiments and edited the manuscript, A.U. refined the PDB coordinates and edited the manuscript, and R.A.B. conceived and supervised the project, and wrote the manuscript.

## Acknowledgements

The authors acknowledge the use of instruments at the Electron Imaging Center for NanoMachines supported by NIH (1S10RR23057, 1S10OD018111 and 1U24GM116792), NSF (DBI-1338135) and CNSI at UCLA. This research was, in part, supported by the National Cancer Institute’s National Cryo-EM Facility at the Frederick National Laboratory for Cancer Research under contract 75N91019D00024. This research was partially funded by The Welch Foundation grant AH-1649.

