## Supplementary Data for "The Hsp60 C-terminus Senses Substrate and Triggers Allosteric ATP Hydrolysis"

### Supplemental

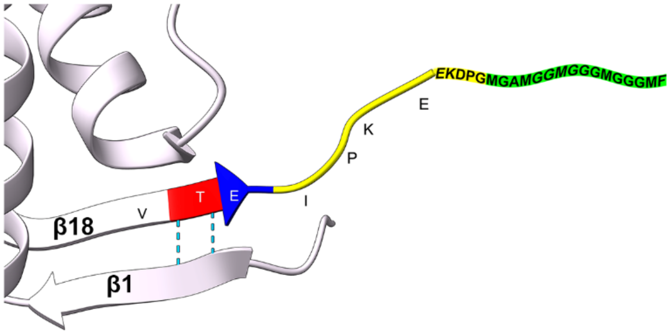


| Protein | C-terminal Tail Truncation | Residues Removed | Assembly of Complex | ATP Hydrolysis |
|---|---|---|---|---|
| Hsp60 | ...VTEIPKEEKDPGMGAMGGMGGGMGGGMF[547] | 0 | tetradecamer | ✓ |
| △532 | ...VTEIPKEEKDPG[531] | 16 | tetradecamer | ✓ |
| △523 | ...VTE[522] | 25 | tetradecamer | × |
| △522 | ...VT[521] | 26 | tetradecamer | × |
| △521 | ...V[520] | 27 | monomer | × |

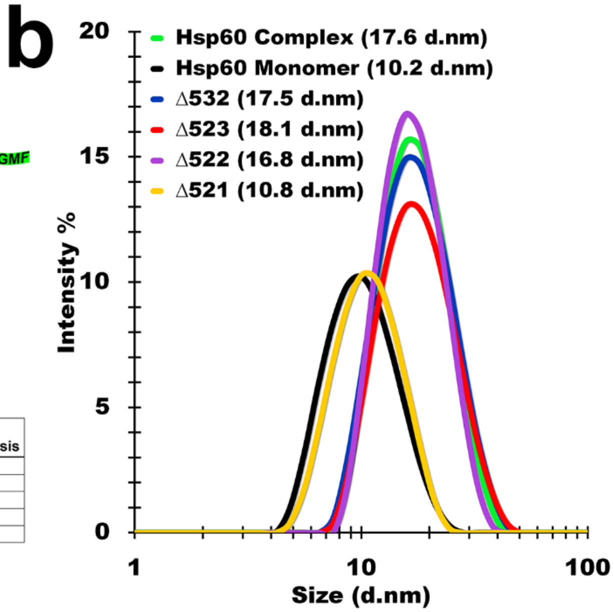


**Supplemental Figure S1.** Physical composition of Hsp60/10 complexes cloned. a) Atlas illustrating how deletion mutants were generated and how they are named. b) Dynamic light scattering (DLS) reveals how purified recombinant proteins were assembled into a tetradecamer or remained as a monomer. The assembled complex of wild-type Hsp60 as well as the monomer of Hsp60 were used as size controls.

#### Secondary Structure

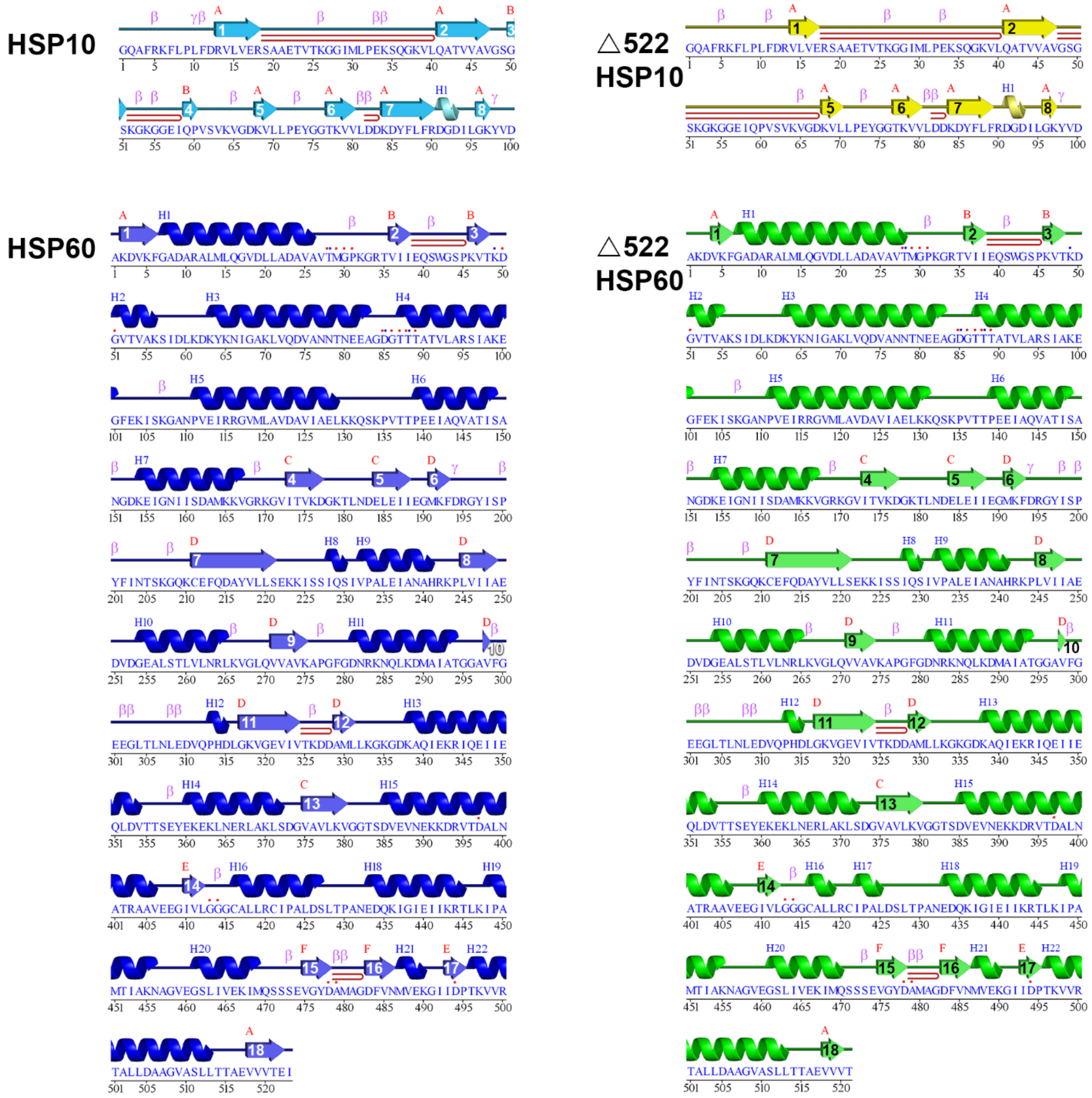


**Supplemental Figure S2**. Secondary structure assignment based on the geometry of the atomic coordinates for Hsp60, Δ522, and the Hsp10 from each reconstruction. In each panel β strands are designated as arrows and numbered inside each arrow, β sheets are assigned a sequential red letter above each strand, and $\alpha$ helices are designated by a colored helix that is numbered in blue above each helix according to the order that they appear in the sequence.

**Supplemental Video S1-** Mechanism of Hsp60-10 ATP hydrolysis

**Supplemental Table S1.** Summary of cryo-EM data collection parameters and resulting quality of model.

| | **Hsp60/10 (Wild type)** | **Hsp60-Δ522/10 (Deletion mutant)** |
|---|---|---|
| Camera | K3 (counting mode) | K3 (counting mode) |
| Magnification | 81,000 | 81,000 |
| Voltage (kV) | 300 | 300 |
| Dose ($e^-/Å^2$) | 40 | 30 |
| Pixel size (Å) | 1.11 | 1.1 |
| Symmetry | D7 | D7 |
| Micrographs collected | 6400 | 7929 |
| Particles picked | 586,721 | 1,145,473 |
| Particles in final reconstruction | 548,970 | 453,877 |
| Reconstruction resolution | 2.3 | 2.6 |
| FSC threshold | 0.143 | 0.143 |
| Deposited PDB code | 8U4M | 8U4G |
| Deposited EMDB map code | EMD-41887 | EMD-41881 |
| Model Composition | | |
| Non-Hydrogen atoms | 68614 | 68460 |
| Protein residues | 8722 | 8694 |
| ATP ligands | 14 | 14 |
| $Mg^{2+}$ ligands | 28 | 28 |
| Waters | 0 | 84 |
| Ramachandran Plot | | |
| Most Favored | 93.0 % | 94.4 % |
| Additionally Allowed | 7.0 % | 5.6 % |
| Disallowed | 0.0 % | 0.0 % |
